# Megakaryocyte TGFβ1 Partitions Hematopoiesis into Immature Progenitor/Stem Cells and Maturing Precursors

**DOI:** 10.1101/689901

**Authors:** Silvana Di Giandomenico, Pouneh Kermani, Nicole Molle, Maria Mia Yabut, Ghaith Abu Zeinah, Thomas Stephens, Nassima Messali, Shahin Rafii, Joseph M. Scandura

## Abstract

Erythropoiesis is a multiweek program coupling massive proliferation with progressive cellular differentiation ultimately enabling a limited number of hematopoietic stem cells (HSCs) to yield millions of erythrocytes per second^1^. Erythropoietin (Epo) is essential for red blood cell (RBC) production but this cytokine acts well after irreversible commitment of hematopoietic progenitor cells (HPCs) to an erythroid fate. It is not known if terminal erythropoiesis is tethered to the pool of available immature hematopoietic stem and progenitor cells (HSPCs). We now report that megakaryocyte-derived TGFβ1 compartmentalizes hematopoiesis by coupling HPC numbers to production of mature erythrocytes. Genetic deletion of TGFβ1 specifically in megakaryocytes (*TGFβ1*^ΔMk/ΔMk^) increased functional HSPCs including committed erythroid progenitors, yet total bone marrow and spleen cellularity and peripheral blood cell counts were entirely normal. Instead, excess erythroid precursors underwent apoptosis, predominantly those erythroblasts expressing the Epo receptor (Epor) but not Kit. Despite there being no deficiency of plasma Epo in *TGFβ1*^ΔMk/ΔMk^ mice, exogenous Epo rescued survival of excess erythroid precursors and triggered exuberant erythropoiesis. In contrast, exogenous TGFβ1 caused anemia and failed to rescue erythroid apoptosis despite its ability to restore downstream TGFβ-mediated Smad2/3 phosphorylation in HSPCs. Thus, megakaryocytic TGFβ1 regulates the size of the pool of immature HSPCs and in so doing, improves the efficiency of erythropoiesis by governing the feed of lineage-committed erythroid progenitors whose fate is decided by extramedullary renal Epo-producing cells sensing the need for new RBCs. Independent manipulation of distinct immature Epo-unresponsive HSPCs within the hematopoietic compartments offers a new strategy to overcome chronic anemias or possibly other cytopenias.

## Background

It is estimated that a third of the world population suffers from anemia^2,3^. Daily production of >200 billion erythrocytes is required to keep up with routine losses so even minor disequilibrium between blood loss and erythrocyte production can lead to anemia. Although many transient anemias are easily treated, therapy for chronic anemias is limited. RBC transfusions and Erythropoiesis Stimulating Agents (ESA) are not always effective yet are linked to significant expense, inconvenience, potential toxicity, and generally transient utility^4,5^. Despite these limitations, development of new approaches to treat chronic anemia have been limited by our incomplete understanding of erythropoiesis.

A great deal of what we know about erythropoiesis relates to the terminal maturation of erythroid precursors leading to hemoglobinization, enucleation and release of immature RBCs into the circulation^6,7^. In the setting of anemia or otherwise reduced oxygen delivery, juxtaglomerular renal Epo producing (REP) cells can increase Epo secretion many thousands-fold thereby serving as a feedback signal reporting erythrocyte need to the marrow^8-13^. Nonetheless, Epo acts during a very narrow window of erythropoiesis, well after progenitor commitment to an exclusively erythroid fate^6,8,9^. It is not known if these final steps of RBC maturation are coupled to the earlier stages of HSPC differentiation; a process that begins almost three weeks earlier when a HSC starts its march towards committed RBC precursors via a series of branching cell fate decisions^7,14,15^.

## Results

### Conditional deletion of TGFβ1 in megakaryocytes (MKs) increases bone marrow apoptosis

Many HSCs reside in close proximity to megakaryocytes where they can be maintained in a quiescent state by niche signals such as Tgfβ1 and Cxcl4^16,17^. Although genetic ablation of Mks using an inducible diphtheria toxin receptor (iDTR) under control of Mk-specific *Cxcl4-cre* driver releases HSCs from quiescence and licenses expansion of the HSC pool, it is not known how this affects blood cell production or other aspects of hematopoiesis. We selectively deleted TGFβ1 in megakaryocytes (*TGFβ1*^ΔMk/ΔMk^) and found that peripheral blood counts were entirely normal in *TGFβ1*^ΔMk/ΔMk^ mice compared to *TGFβ1*^FL/FL^ littermate controls despite the pool of primitive hematopoietic cells being expanded (Fig. 1a). Sequestration of maturing cells within hematopoietic tissues could not explain this discrepancy because total bone marrow and spleen cellularity was normal in *TGFβ1*^ΔMk/ΔMk^ mice (Fig. 1b). Excess HSCs in *TGFβ1*^ΔMk/ΔMk^ mice appeared capable of robust differentiation because the number of immature lineage-negative (Lin^neg^) hematopoietic progenitor cells was increased in the marrows of *TGFβ1*^ΔMk/ΔMk^ mice (Fig. 1c). Thus, it remained unexplained why the expanded number of HSCs and HPCs do not increase blood counts and marrow cellularity.

**Figure 1.**
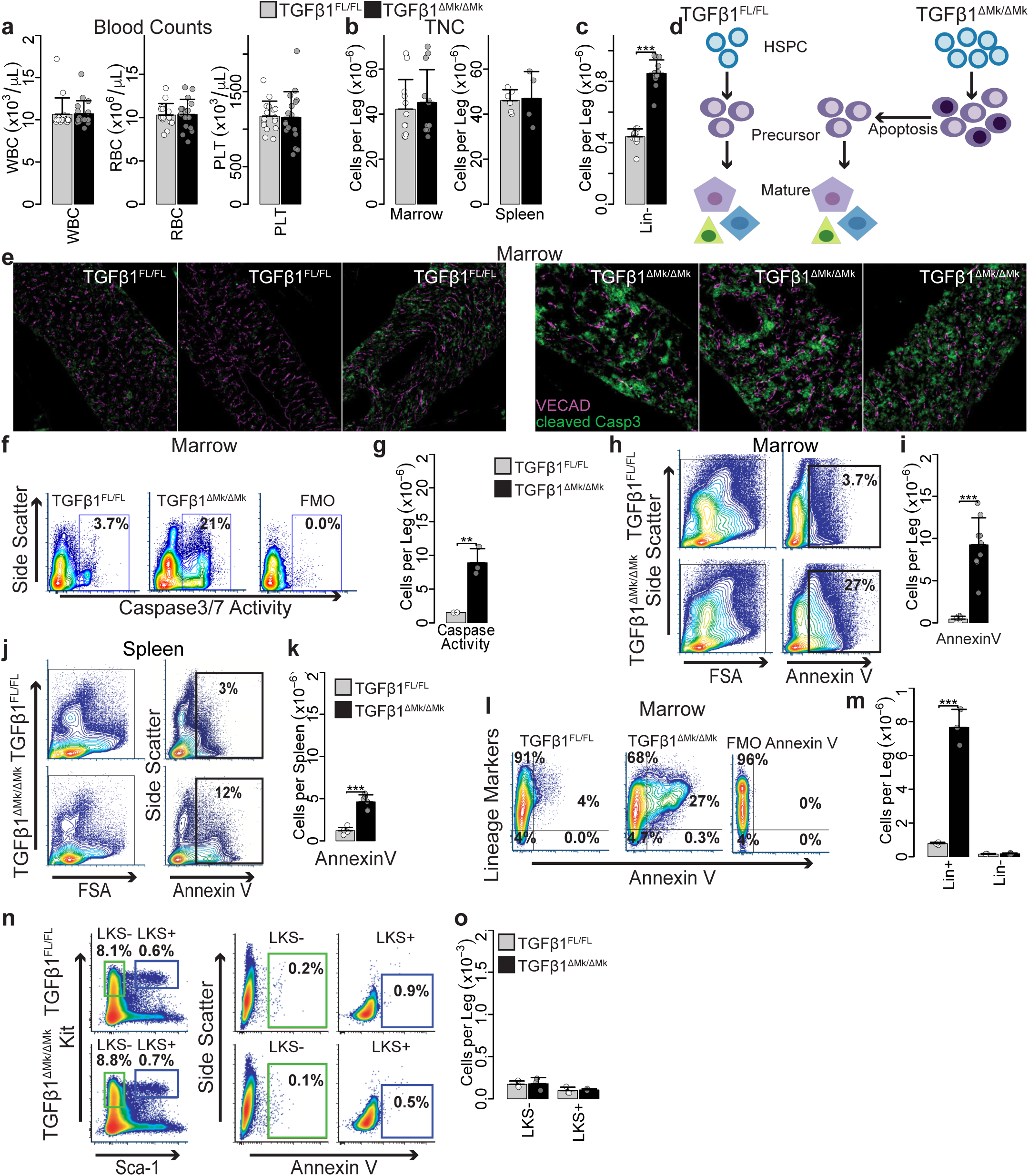
Conditional deletion of *TGFβ1* in megakaryocytes (MKs) increases apoptosis. **a**, White blood cell (WBC), RBC, and platelet counts are shown for *TGFβ1*^FL/FL^ and *TGFβ1*^ΔMk/ΔMk^ mice (n=16 per group). **b**, Whole BM (Marrow) (n=12 per group) and spleen (n=5 per group) cellularity are shown for *TGFβ1*^FL/FL^ (grey) and *TGFβ1*^ΔMk/ΔMk^ mice (black). **c**, The number of Lin^neg^ (those not expressing mature lineage markers CD3, B220, CD11b, Gr1 or Ter119) marrow cells per leg (femur & tibia) is shown (n=13 per group). **d**, Schematic of model for how excess immature HSPCs may yield normal marrow cellularity and blood cell counts due to apoptosis of surplus hematopoietic precursors. **e**, Confocal immunofluorescence imaging of cleaved Caspase3 (green) and Cdh5 (magenta) in bone marrow sections of *TGFβ1*^FL/FL^ and *TGFβ1*^ΔMk/ΔMk^ mice (n=3 per group). **f**, Representative flow cytometry data of cleaved Caspase3/7 activity is shown for *TGFβ1*^FL/FL^ and *TGFβ1*^ΔMk/ΔMk^ mice and for the fluorescence minus one (FMO) negative control. **g**, Quantification of apoptotic cells as assessed using the Caspase3/7 reporter (n=3 per group). **h**, Representative flow cytometry data showing AnnexinV staining in marrow from TGFβ1^FL/FL^ (top row) and *TGFβ1*^ΔMk/ΔMk^ (bottom row) mice. **i**, Quantification of the number of AnnexinV^+^ apoptotic cells in the marrow of *TGFβ1*^FL/FL^ and *TGFβ1*^ΔMk/ΔMk^ mice (n=9 per group). **j**, Representative data for spleen, as shown for marrow in panel h. **k**, Quantification of the number of AnnexinV^+^ apoptotic cells in the spleen of *TGFβ1*^FL/FL^ and *TGFβ1*^ΔMk/ΔMk^ mice (n=6 per group). **l**, Representative flow cytometry data showing lineage markers (Lin=CD3, B220, CD11b, Gr1 or Ter119) plotted against AnnexinV in marrow from *TGFβ1*^FL/FL^ and *TGFβ1*^ΔMk/ΔMk^ mice. **m**, Quantification of the number of AnnexinV^+^ apoptotic Lin+ and Lin^neg^ cells in the marrow of *TGFβ1*^FL/FL^ and *TGFβ1*^ΔMk/ΔMk^ mice (n=8). **n**, Gating and representative flow cytometry data showing AnnexinV staining in LKS^neg^ and LKS^+^ marrow populations for *TGFβ1*^FL/FL^ (top row) and *TGFβ1*^ΔMk/ΔMk^ (bottom row) mice. **o**, Quantification of the number of AnnexinV^+^ apoptotic LKS^neg^ and LKS^+^ cells in the marrow of *TGFβ1*^FL/FL^ and *TGFβ1*^ΔMk/ΔMk^ mice (n=3 per group). All the quantified data are shown as mean ± SEM (* P<0.05, ** P<0.01, *** P<0.001, or if not shown, the comparison was not significant).

Hematopoietic cell population size is determined by the balance of cell gain (proliferation/self-renewal and differentiation) and cell loss (apoptosis and differentiation). Many late acting hematopoietic cytokines are not required for lineage commitment yet provide essential proliferation, differentiation and survival signals during maturation of hematopoietic precursors^6,18^. Thus, we hypothesized that the excess progenitors observed in the *TGFβ1*^ΔMk/ΔMk^ mice failed to increase blood counts because their progeny were unneeded, and inadequately supported by homeostatic levels of late-acting cytokines (Fig. 1d). Indeed, bone marrow apoptosis was markedly increased in the *TGFβ1*^ΔMk/ΔMk^ mice compared to controls, as reported by cleaved caspase 3 (Fig. 1e-g and Supplementary Fig. 1a-c) or AnnexinV binding (Fig. 1h-k). The excess apoptotic cells seemed largely restricted to hematopoietic cells expressing the lineage markers (Lin) Gr1, CD11b, CD3, B220, or Ter119 (Fig. 1l-m). Annexin V staining of lineage-marker negative (Lin^neg^), Kit^+^ Sca1^neg^ (LKS^neg^) HPCs and LKS^+^ HSPCs was rare in both *TGFβ1*^*ΔMk/ΔMk*^ mice and littermate controls (Fig. 1n-o). These results support an interpretation that excess, unneeded hematopoietic precursors are pruned by apoptosis during hematopoietic differentiation.

### Increased hematopoietic stem and progenitor cells in TGFβ1^ΔMk/ΔMk^ mice have normal function

To rule out the possibility that progeny of HSPCs from *TGFβ1*^*ΔMk/ΔMk*^ mice are intrinsically defective, we performed competitive repopulation assays by co-transplanting bone marrow from *TGFβ1*^ΔMk/ΔMk^ mice or *TGFβ1*^FL/FL^ littermates competitively with CD45.1 congenic donor cells. As expected, *TGFβ1*^FL/FL^ donor cells engrafted and contributed to multi-lineage hematopoiesis indistinguishably from the congenic CD45.1 controls (Fig. 2a and Supplementary Fig. 1d-f). In contrast, *TGFβ1*^*ΔMk/ΔMk*^ donor cells outcompeted control cells confirming that *TGFβ1*^*ΔMk/ΔMk*^ donors are enriched for functional HSCs and indicating that these *TGFβ1*^*ΔMk/ΔMk*^ HSCs yield progeny fully capable of reconstituting hematopoiesis after transplant. Serial transplantation of recipients demonstrated HSC self-renewal and contribution to hematopoiesis was intrinsically normal in the *TGFβ1*^ΔMk/ΔMk^ donor cells (Fig. 2b and Supplementary Fig. 1g-h).

**Figure 2.**
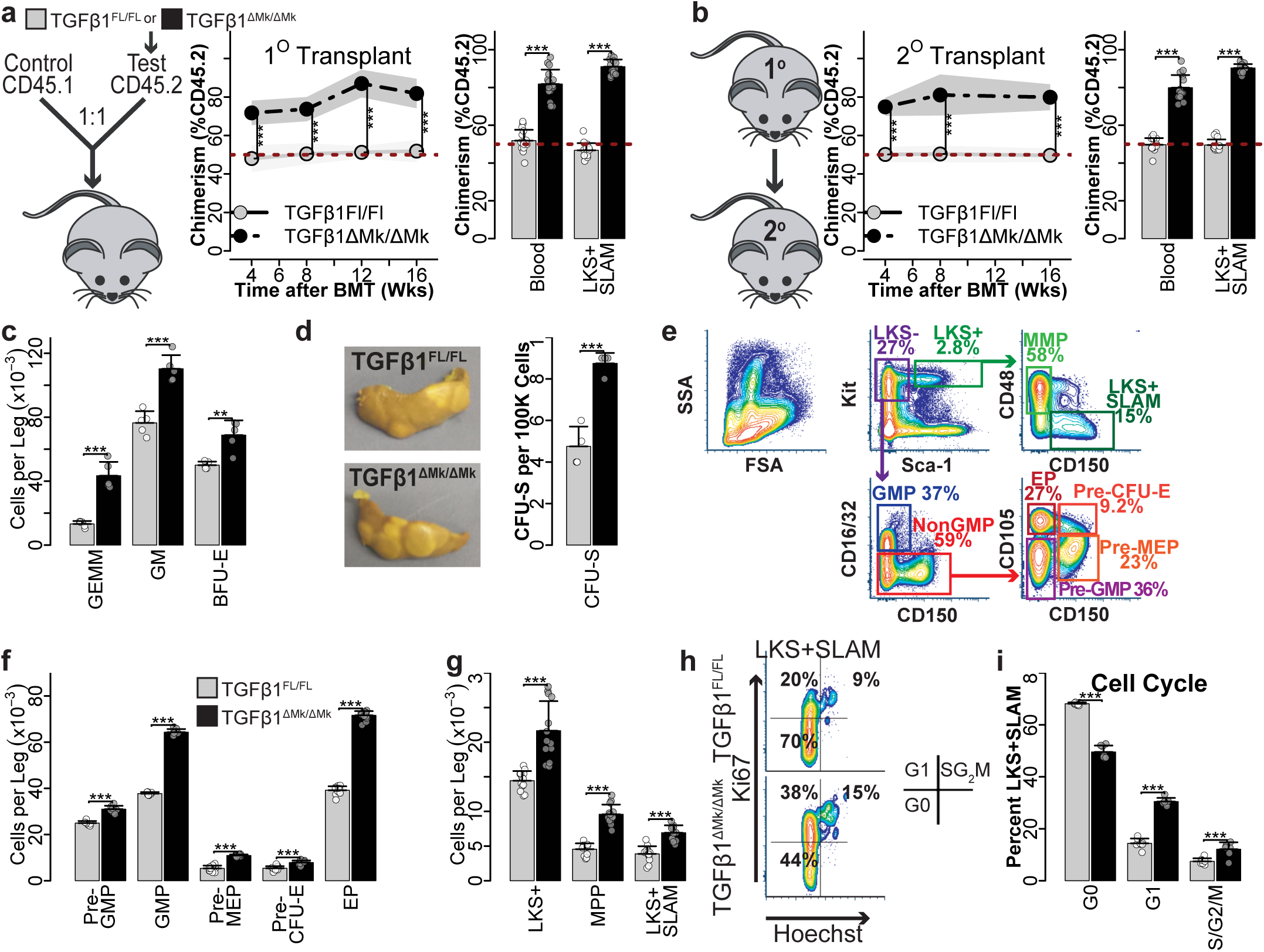
Increased hematopoietic stem and progenitor cells (HSPCs) in *TGFβ1*^ΔMk/ΔMk^ mice have normal function. **a**, Schematic for the competitive repopulation assay is shown. Marrow CD45.2 donor cells from either from *TGFβ1*^FL/FL^ or *TGFβ1*^ΔMk/ΔMk^ mice were mixed in a 1:1 ratio with marrow donor cells from congenic CD45.1 mice, then transplanted into lethally irradiated mice (n=8 per group). CD45.2 chimerism in peripheral blood is shown at the indicated times after transplant (left). Chimerism of blood and bone marrow LKS+SLAM (Lin^neg^, Kit^+^, Sca1^+^, CD150^+^, CD48^neg^) cells is shown 16 weeks after transplant for mice having received *TGFβ1*^FL/FL^ (grey) or *TGFβ1*^ΔMk/ΔMk^ (black) donor cell grafts, as assessed by flow cytometry. **b**, Schematic for secondary transplantation of marrow harvested from primary recipients is shown. CD45.2 chimerism in peripheral blood is shown at the indicated times after secondary transplant (left). Chimerism in blood and in peripheral blood and marrow LKS+SLAM is shown 16 weeks after secondary transplant as for panel a (n=8 per group). **c**, CFC enumeration of functional myeloid HPCs is shown for *TGFβ1*^FL/FL^ and *TGFβ1*^ΔMk/ΔMk^ marrow cells (n=5 per group). The total number of CFU-GEMM (GEMM), CFU-GM (GM) and BFU-E are shown per leg (femur & tibia). **d**, The number of spleen colony forming units (CFU-S_12_) 12 days after transplantation is shown per 100,000 donor cells transplanted (n=4 per group). **e**, Gating and representative flow cytometry data is shown for the indicated HSPC populations. **f**, Quantification of the number of marrow cells with the immunophenotype of CMP, GMP, MEP, EP, Pre-CFU-GM, Pre-CFU-GMP, Pre-MEP is shown for *TGFβ1*^FL/FL^ and *TGFβ1*^ΔMk/ΔMk^ mice (n=12 per group). **g**, The number of LKS^+^ (Lin^neg^Kit^+^Sca1^+^), MPP and LKS+SLAM cells per leg is shown for *TGFβ1*^FL/FL^ and *TGFβ1*^ΔMk/ΔMk^ mice (n=12 per group). **h**, Gating and representative flow cytometry data is shown for LKS+SLAM cell cycle as reported by Ki67 and Hoechst staining. **i**, Cell cycle of BM LKS+SLAM HSCs is shown for *TGFβ1*^FL/FL^ and *TGFβ1*^ΔMk/ΔMk^ (mice n=6 per group). All the quantified data are shown as mean ± SEM (* P<0.05, ** P<0.01, *** P<0.001, or if not shown, the comparison was not significant).

*Ex vivo* HPC function appeared normal in *TGFβ1*^*ΔMk/ΔMk*^ mice suggesting that these cells were not intrinsically defective in *TGFβ1*^*ΔMk/ΔMk*^ mice (Fig. 2c). The number of granulocytic-monocytic progenitors (GMP), burst forming unit erythroid progenitors (BFU-E) and granulocyte, erythroid, monocyte, megakaryocyte progenitors (CFU-GEMM) were all increased in the marrow of *TGFβ1*^*ΔMk/ΔMk*^ mice as reported by colony forming cell (CFC) assays. Similarly, the 12-day colony forming unit spleen (CFU-S_12_) assay showed a greater number of functionally normal MPPs in *TGFβ1*^*ΔMk/ΔMk*^ donor marrow compared to *TGFβ1*^FL/FL^ littermate controls (Fig. 2d). Quantified HSPC immunophenotypes correlated well with functional measures again demonstrating that the number of HSPCs is increased in *TGFβ1*^*ΔMk/ΔMk*^ mice (Fig. 2e-g). The larger number of HSPCs and their more proliferative character in *TGFβ1*^*ΔMk/ΔMk*^ mice (Fig. 2h-i) indicate that megakaryocytic TGFβ1 normally curbs the HSPC pool size. Yet progeny of these excess HSPCs appear to undergo apoptosis rather than contribute to bone marrow cellularity and mature blood cell counts.

### Excess erythroid precursors undergo apoptosis in vivo

The islands of apoptotic cells observed in bone marrow of *TGFβ1*^ΔMk/ΔMk^ mice (Fig. 1e) are reminiscent of erythroid islands suggesting that excess erythroid precursors may be contributing to the formation of these clusters. Indeed, the proportion of Annexin V-staining apoptotic Ter119^+^ erythroid precursor cells (EPCs) was ∼6-fold higher in the marrow and spleen of *TGFβ1*^ΔMk/ΔMk^ mice compared to *TGFβ1*^FL/FL^ littermate controls (Fig. 3a-b). We quantified apoptosis within well-defined populations of maturing erythroid precursors^19-21^ (Fig. 3c) using Annexin V. Although the number of immature erythroid precursors—such as pro/basophilic (EI), polychromatophilic (EII), and orthochromatophilic erythroblasts (EIII)—were increased in *TGFβ1*^ΔMk/ΔMk^ mice, the excess seemed largely comprised of cells staining for the apoptotic marker Annexin V (Fig. 3d). In contrast, the number of mature reticulocytes and erythrocytes (EIV/EV) in *TGFβ1*^ΔMk/ΔMk^ mice was indistinguishable from *TGFβ1*^FL/FL^. Thus, the excess erythroid committed precursors are culled during early maturation thereby producing normal numbers of erythrocytes and normal red blood cell counts (RBCs).

**Figure 3.**
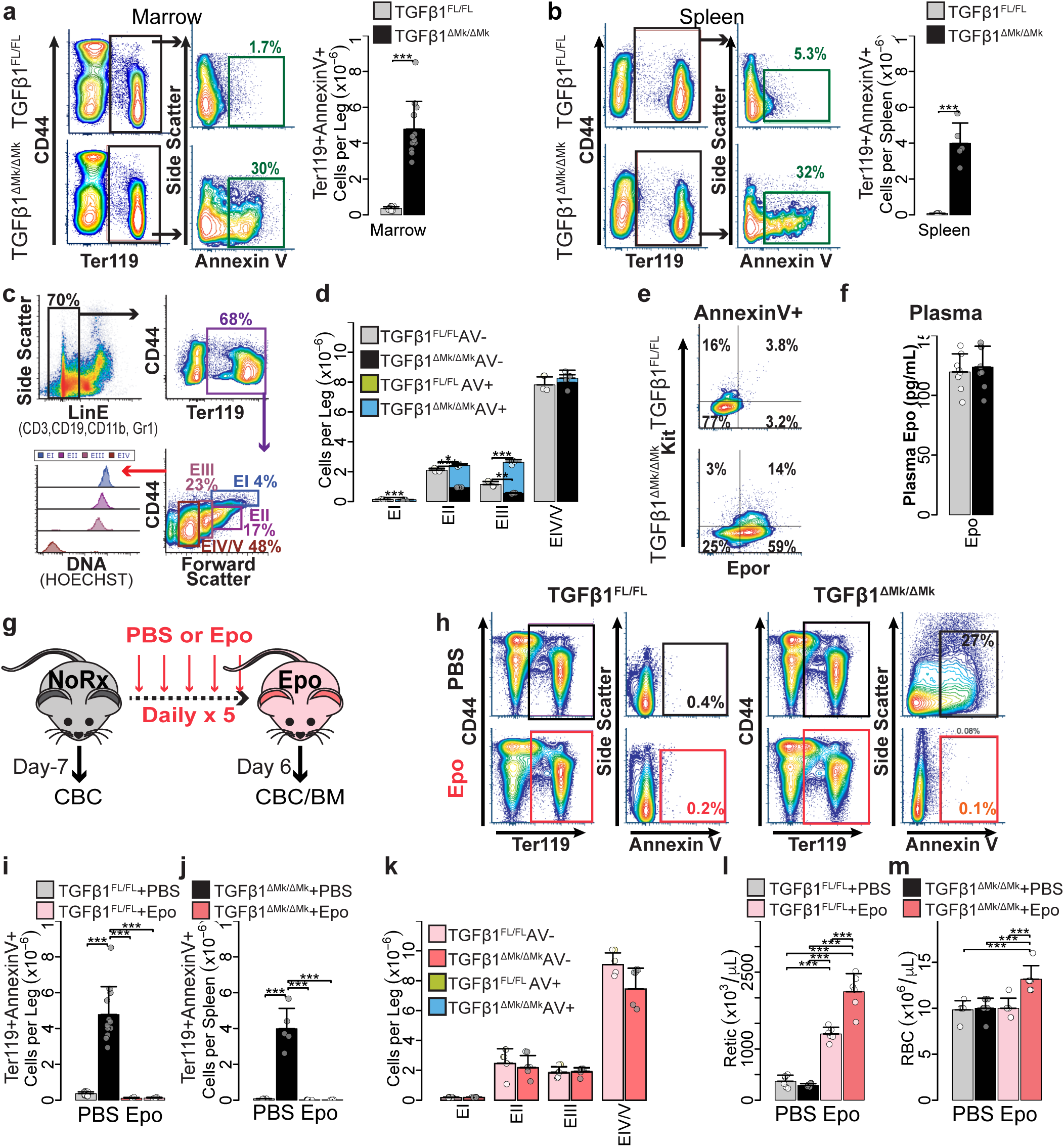
Surplus Epo-dependent erythroid precursors undergo apoptosis *in vivo.* **a**, Gating and representative flow cytometry data is shown for AnnexinV staining of Ter119^+^ erythroid precursors in marrow from *TGFβ1*^FL/FL^ (top row) and *TGFβ1*^ΔMk/ΔMk^ (bottom row) mice. Quantification of the number of marrow, AnnexinV-staining, apoptotic Ter119^+^ erythroid cells is shown for *TGFβ1*^FL/FL^ and *TGFβ1*^ΔMk/ΔMk^ mice (n=12 per group). **b**, Gating and representative flow cytometry data is shown spleen cells as described for panel a. Quantification of splenic apoptotic Ter119^+^ erythroid cells is shown for *TGFβ1*^FL/FL^ and *TGFβ1*^ΔMk/ΔMk^ mice (n=4 to 6 per group). **c**, Flow cytometry gating strategy using LinE- (CD3, B220, Gr1, CD11b), CD44, Ter119, Hoechst (DNA) and AnnexinV to characterize erythroid maturation. **d**, Quantification of erythroid precursors is shown (n=3 per group). Viable cells and AnnexinV^+^ apoptotic cells for *TGFβ1*^FL/FL^ (grey/green) and *TGFβ1*^ΔMk/ΔMk^ (black/blue) mice, respectively. **e**, Representative flow cytometry showing staining of Kit and Epor within the AnnexinV^+^/Ter119^+^ population identifies excess Epor+/Kit- erythroblasts within the *TGFβ1*^ΔMk/ΔMk^ marrow. **f**, Plasma EPO levels are shown as assessed by ELISA (n=8 per group). **g**, *TGFβ1*^FL/FL^ and *TGFβ1*^ΔMk/ΔMk^ mice were treated with 300U/kg Epo daily for 5 days and sacrificed on day 6 for analysis. **h**, Representative AnnexinV staining of marrow erythroid precursor cells is shown after PBS (top) or Epo (bottom) treatment for *TGFβ1*^FL/FL^ (left) and *TGFβ1*^ΔMk/ΔMk^ (right) mice. The number of AnnexinV^+^ apoptotic Ter119^+^ erythroid precursor cells is shown in marrow (**i**) and spleen (**j**) before and after Epo treatment (n=4 to 12 per group). **k**, Apoptosis within maturing erythroid precursors is shown for marrow cells after Epo treatment, as shown in panel d. Viable cells and AnnexinV^+^ apoptotic cells for Epo-treated *TGFβ1*^FL/FL^ (pink/green) and *TGFβ1*^ΔMk/ΔMk^ (red/blue) mice, respectively (n=4 per group). Reticulocyte counts (**l**) and RBCs (**m**) are shown for *TGFβ1*^FL/FL^ (grey/pink) and *TGFβ1*^ΔMk/ΔMk^ (black/red) mice before and after Epo (n=6 per group). All the quantified data are shown as mean ± SEM (* P<0.05, ** P<0.01, *** P<0.001, or if not shown, the comparison was not significant).

### Excess erythroid precursors are rescued by exogenous Epo

Apoptotic Ter119^+^ erythroid precursors within *TGFβ1*^ΔMk/ΔMk^ marrow predominantly expressed the erythropoietin (Epo) receptor (Epor) but not Kit (Fig. 3e). Epo provides a survival signal to Epor+ erythroid precursors allowing them to escape apoptosis and continue differentiation^8-10^. Excess apoptosis of erythroid precursors in *TGFβ1*^ΔMk/ΔMk^ mice was not due to subnormal plasma Epo levels (Fig. 3f). We reasoned that the surplus, unneeded Epor+ cells may not be supported by physiologic Epo levels. To test this, we treated mice with exogenous Epo (300 U/kg) (Fig. 3g). Strikingly, exogenous Epo rescued the excess apoptosis of erythroid precursors in *TGFβ1*^ΔMk/ΔMk^ marrow and spleen (Fig. 3h-k). The erythropoietic response of *TGFβ1*^ΔMk/ΔMk^ mice to Epo was much more robust compared to *TGFβ1*^FL/FL^ littermate controls (Fig. 3l>-m) suggesting that the rescued apoptosis resulted in increased erythropoietic output. Using phenylhydrazine (PHZ) induced hemolysis as an anemia model, we found that the erythropoietic response of *TGFβ1*^ΔMk/ΔMk^ mice was much brisker and more robust than that of *TGFβ1*^FL/FL^ littermate controls (Supplementary Fig. 2). These results demonstrate that although genetic deletion of *TGFβ1* in megakaryocytes expands the number of committed erythroid progenitors, the resultant glut of their erythroid precursor progeny requires non-homeostatic Epo levels to promote survival, expansion and maturation to erythrocytes.

### The dominant source of TGFβ1 signaling in HSPCs is produced by Megakaryocytes

Megakaryocytic TGFβ1 is necessary for robust phosphorylation of receptor activated Smad2 and Smad3 (Smad2/3) in HSPCs, as assessed by flow cytometry with intracellular staining (Fig. 4a). In contrast, we detected little phospho-Smad2/3 (pSmad2/3) in erythroid precursors in either the *TGFβ1*^ΔMk/ΔMk^ mice or *TGFβ1*^FL/FL^ littermate controls (Fig. 4a). Pretreating mice with a TGFβ1 neutralizing antibody (1D11) reduced the mean fluorescence intensity (MFI) of pSmad2/3 in HSPCs compared to an isotype control (13C4) antibody confirming the specificity of the phospho-flow cytometry assay (Supplementary Fig. 3). HSPCs in *TGFβ1*^ΔMk/ΔMk^ mice were capable of normal TGFβ1 signaling because exogenous TGFβ1 induced strong phosphorylation of Smad2/3, restoring levels to those in *TGFβ1*^FL/FL^ controls (Fig. 4b-c). Thus, megakaryocytes serve as the major source of TGFβ triggering Smad2/3 phosphorylation in HSPCs.

**Figure 4.**
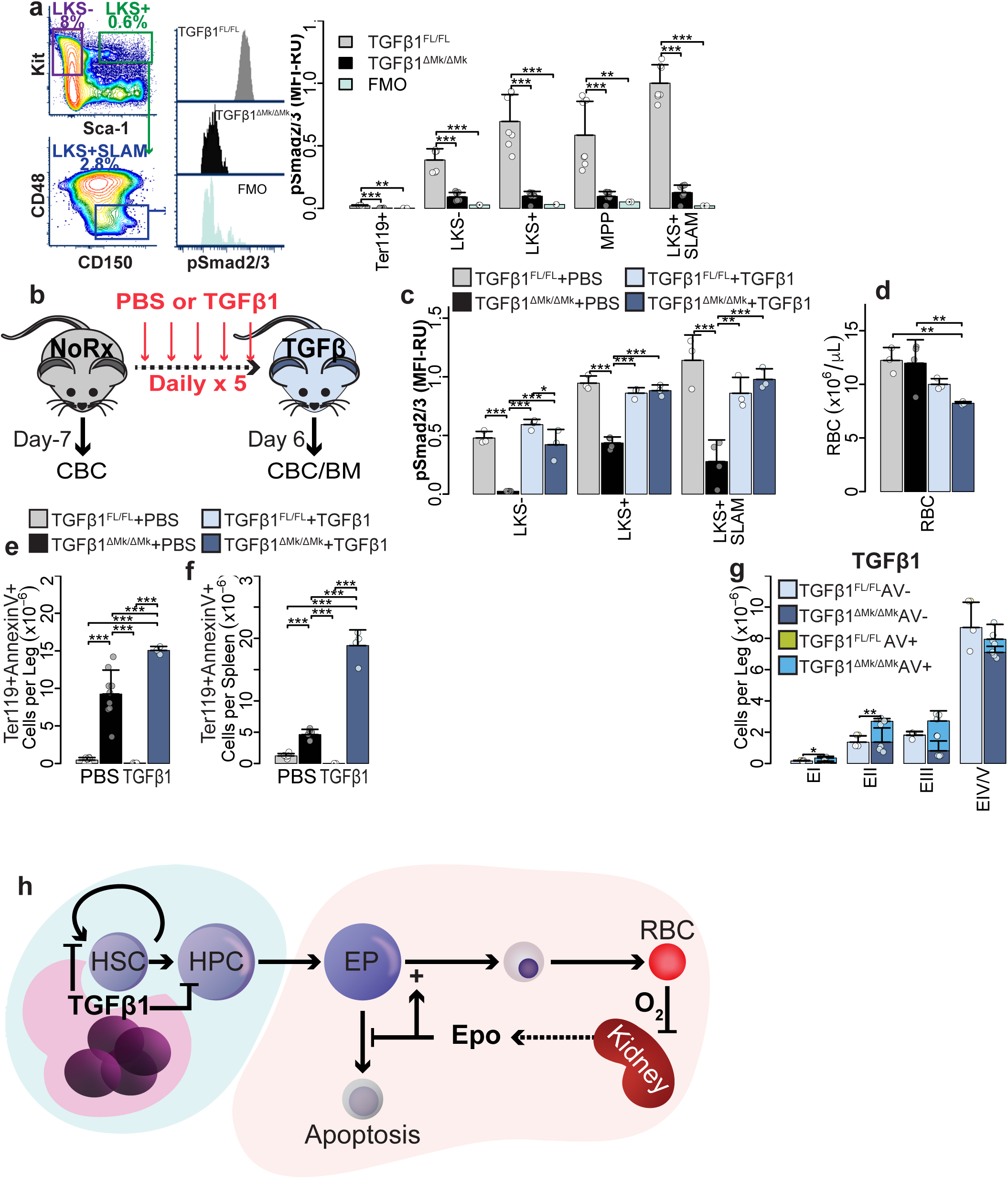
Megakaryocytes direct TGFβ1 signaling in HSPCs but erythroid precursor maturation is independently regulated. **a**, Gating and representative flow cytometry data assessing Smad2/3 phosphorylation (pSmad2/3) within the LKS+SLAM population is shown for *TGFβ1*^FL/FL^ and *TGFβ1*^ΔMk/ΔMk^ mice and for the FMO negative control. Mean fluorescence intensity (MFI) of pSmad2/3 within the indicated hematopoietic populations is shown for *TGFβ1*^FL/FL^ and *TGFβ1*^ΔMk/ΔMk^ mice (n=7 per group). **b**, Schematic showing *TGFβ1*^FL/FL^ and *TGFβ1*^ΔMk/ΔMk^ mice treated with 5μg/kg TGFβ1 daily for 5 days and sacrificed on day 6 for analysis. **c**, MFI of pSmad2/3 staining is shown for Lin^neg^ HSPC subpopulations as quantified by flow cytometry of *TGFβ1*^FL/FL^ (grey/light blue) and *TGFβ1*^ΔMk/ΔMk^ (black/dark blue) mice treated with PBS or TGFβ1 (n=3 to 4 mice per group). **d**, RBC counts are shown for *TGFβ1*^FL/FL^ and *TGFβ1*^ΔMk/ΔMk^ mice treated with PBS or TGFβ1 (n= 6 to 7 per group). The number of Ter119^+^AnnexinV^+^ apoptotic cells in marrow (**e**) and spleen (**f**), is shown for *TGFβ1*^FL/FL^ or *TGFβ1*^ΔMk/ΔMk^ mice (n=4 to 9 per group). **g**, Apoptosis within maturing erythroid precursors is shown for marrow cells after TGFβ1 treatment (n=3 to 4 per group), as shown in panel 3d. **h**, Schematic of proposed compartmentalized model of erythropoiesis wherein megakaryocytic TGFβ1 serve as a gatekeeper regulating the feed of committed erythroid progenitors to a maturation module regulated by Epo thereby coupling erythroblast survival to the need for RBC production as sensed by renal Epo-producing cells. All the quantified data are shown as mean ± SEM (* P<0.05, ** P<0.01, *** P<0.001, or if not shown, the comparison was not significant).

### Exogenous TGFβ1 cannot rescue the erythroid phenotype of TGFβ1^ΔMk/ΔMk^ mice

It is possible that TGFβ1 sensitizes erythroid precursors to Epo, and if true, exogenous TGFβ1 should rescue erythroid precursor dropout in *TGFβ1*^ΔMk/ΔMk^ mice. To test this, we treated the mice with TGFβ1 for 5 days and assessed erythroid response (Fig. 4b). Although exogenous TGFβ1 reestablished pSmad2/3 in *TGFβ1*^ΔMk/ΔMk^ HSPCs (Fig. 4c), it did not trigger an erythropoietic response. Rather, exogenous TGFβ1 induced mild anemia in the *TGFβ1*^ΔMk/ΔMk^ mice (Fig. 4d) coupled with worsened apoptosis in the marrow and spleen (Fig. 4e-g). In contrast, exogenous TGFβ1 did not induce detectable changes in RBCs or apoptosis in *TGFβ1*^FL/FL^ controls (Fig. 4e-g). Therefore, the boundary of megakaryocytic TGFβ1 activity is compartmentalized within the marrow with predominant effects on immature HSPCs while excluding their progeny (Fig. 4h).

## Discussion

Erythropoiesis is subject to modular regulation. Epo acts during a very limited stage of differentiation—supporting early erythroid precursors (CFUe/pre-erythroblasts) as they gear up for iron accumulation, heme synthesis and globin gene transcription^7,14,15^. Accordingly, many causes of anemia are unresponsive to exogenous Epo. Recently, TGFβ superfamily activin receptor (Acvr2a/b) ligand traps have shown activity treating chronic Epo-unresponsive anemia in myelodysplastic syndrome (MDS) and beta-thalassemia and are thought to promote erythroid maturation after Epo signaling^22-26^. However, the molecular regulators of the Epo-unresponsive erythroid compartment are not well defined. Here we show that the number of committed erythroid progenitors is controlled by the availability of megakaryocytic TGFβ1; adding a new level of erythroid regulation prior to the Epo restriction point. Thus, megakaryocytic TGFβ1 is a gate-keeper, matching the number of immature hematopoietic progenitor and stem cells to the production of mature effector cells. Megakaryocytes and erythroids share a common progenitor, but megakaryocytes also have a direct ontological link to a subset of Mk-biased HSCs^27^. It will be important to understand the evolutionary reasoning for megakaryocytes being placed at the helm of hematopoiesis and to explore the potential of manipulating this pathway clinically.

## Supplemental Figures

**Supplementary Figure 1. a**, Representative flow cytometry showing intracellular staining for cleaved Caspase3 in mice and the fluorescence minus one (FMO) negative control. **b**, Quantification of apoptotic cells as assessed by intracellular staining for cleaved Caspase3 as shown in panel a. **c**, Slot blot for cleaved Caspase3 in *TGFβ1*^FL/FL^ and *TGFβ1*^ΔMk/ΔMk^. **d**, Schematic for the competitive repopulation assay is shown. WBM CD45.2 donor cells from either from *TGFβ1*^FL/FL^ or *TGFβ1*^ΔMk/ΔMk^ mice were mixed in a 1:1 ratio with WBM donor cells from congenic CD45.1 mice, then transplanted into sub-lethally irradiated mice **e**, Representative flow cytometry showing lymphoid (CD3/CD19) and myeloid (Gr1/CD11b) proportion of CD45.2+ cells in peripheral blood for *TGFβ1*^FL/FL^ (top) or *TGFβ1*^ΔMk/ΔMk^ (bottom) mice is shown. **f**, Quantification of the myeloid and lymphoid proportions is shown for primary transplant recipients of *TGFβ1*^FL/FL^ (grey) or *TGFβ1*^ΔMk/ΔMk^ (black) donors is shown (n=8 per group). **g**, Schematic for secondary transplantation is shown. **h**, Quantification of the myeloid and lymphoid proportions is shown for secondary transplant recipients of *TGFβ1*^FL/FL^ (grey) or *TGFβ1*^ΔMk/ΔMk^ (black) donors is shown (n=8 per group). All the quantified data are shown as mean ± SEM (* P<0.05, ** P<0.01, *** P<0.001, or if not shown, the comparison was not significant.

**Supplementary Figure 2. a**, Schematic showing the schedule for induction of hemolysis with 60 mg/kg/day phenylhydrazine (PHZ) on day 1 and 2 and monitoring of blood count recovery (days 3, 5, 8, 14). Recovery of (**b**) RBCs, (**c**) PLTs and (**d**) WBCs was monitored before and after PHZ induced hemolysis in *TGFβ1*^FL/FL^ and *TGFβ1*^ΔMk/ΔMk^ mice (n=4 per group). Data are shown as mean ± SEM (* P<0.05, ** P<0.01, *** P<0.001, or if not shown, the comparison was not significant.

**Supplementary Figure 3. a**, C57BL/6J mice were treated with either a TGFβ-neutralizing antibody (1D11) or isotype control antibody (13C4) at a dose of 10 mg/kg on days 1, 5 and 10 and then analyzed on day 16. **b**, TGFβ signaling was assessed in LKS^neg^, LKS^+^ and LKS+SLAM cells by flow cytometry with intracellular staining for pSmad2/3. Mean fluorescence intensity (MFI) of pSmad2/3 staining is shown for mice treated with 13C4 (green) or 1D11 (blue) (n=4 per group). Data are shown as mean ± SEM (* P<0.05, ** P<0.01, *** P<0.001, or if not shown, the comparison was not significant.

## Materials and Methods

### Animals

Transgenic C57BL/6 mice with Cre recombinase driven from the regulatory elements of Cxcl4 (Platelet factor 4, Pf4), C57BL/6-Tg(Pf4-cre)Q3Rsko/J (Pf4-cre)^28^, were crossed with mice harboring a floxed *TGFβ1* locus^29^ mice (Tgfb1^tm2.1Doe^, *TGFβ1*^FL/FL^ mice) to generate Pf4-Cre/ *TGFβ1*^FL/FL^ mice with selective deletion of TGFβ1 in megakaryocytes and platelets (*TGFβ1*^ΔMk/ΔMk^ mice). Animals were maintained in the Weill Cornell Medicine Animal Facility. All protocols were approved by the Weill Cornell Medicine Institutional Animal Care and Use Committee.

### Immunophenotypic analysis of BM HSPCs and HSCs by flow cytometry

Mice were killed by CO_2_ asphyxiation and bones (femur, tibia, +/-humeri) were dissected free of muscle and tendons and crushed in DMEM using a mortar and pestle. The resulting cell suspension was filtered through a 40-µm mesh and washed in PEB (2 mM EDTA, 0.2% BSA in PBS, pH 7.4). Spleens were isolated, and then minced before grinding through a 40-µm mesh to generate single-cell suspensions. Lin^−^ cells were purified using a biotinylated lineage cell depletion cocktail (Miltenyi Biotec).

Lineage-depleted cells were stained with PerCP/Cy5.5-CD117 (BD), PE/Cy7-Sca-1 (BioLegend), Alexa Fluor 700–CD48 (BioLegend) and PE-CD150 (BioLegend), APC/Cy7-CD16/32 (FcR) (BioLegend), APC-CD105 (BioLegend). Hematopoietic populations were identified as following: LT-HSCs, LKS^+^CD48^neg^CD150^+^; MPPs, LKS^+^CD48^+^CD150^neg^; granulocyte macrophage progenitors (GMPs), LKS^neg^CD150-FcR-; Pre-GMP, LKS^neg^CD150+CD105^neg^; Pre-megakaryocyte erythroid progenitors (MEPs), LKS^neg^CD150+FcR^neg^CD105^neg^; Pre-CFU-Erythroid (Pre-CFU-E) LKS^neg^CD150+FcR^neg^CD105+; Erythroid Progenitors (EP) LKS^neg^CD150^neg^FcR^neg^CD105+^30^

To analyze cell cycle or pSmad2/3 staining in HSPCs, lineage-depleted cells were stained with streptavidin (BioLegend), APC-CD117 (BD), and PE/Cy7-Sca-1 (BioLegend), Alexa Fluor 700–CD48 (BioLegend) and PE or BV605-CD150 (BioLegend) and then fixed and permeabilized using CytoFix/Perm (BD). For cell cycle, fixed permeabilized cells were stained with PerCP/Cy5.5-Ki67 (BD) and Hoechst 33342 (Invitrogen). To analyze pSmad2/3 signal, fixed permeabilized cells were stained first with pSmad2/3 primary antibody (Cell Signaling Technology) and then with secondary anti-rabbit, antibody Cy3 (Jackson Labs).

Specificity of pSmad2/3 staining was tested by treating C57BL/6J mice with either the TGFβ-neutralizing antibody 1D11 (10 mg/kg) or a non-targeting control antibody 13C4 (10 mg/kg) on day 5, 10 and 15. Mice were sacrificed on day 16 for analysis. Smad2/3 phosphorylation was assessed by intracellular flow cytometry as described previously.

Analysis gates were set based upon the fluorescence minus one (FMO) fluorophore. Multidimensional FACS analysis was performed using a BD LSRII equipped with five lasers (BD). FlowJo (Tree Star) and FCS Express (De Novo) were used to analyze flow cytometry data.

### Immunophenotypic analysis of erythroid precursors by flow cytometry

Mice were killed by CO_2_ asphyxiation and bones or spleen were processed as above except lineage depletion was not performed. Cell suspensions were stained with APC-conjugated Ter119, APC/Cy7 conjugated CD44, Hoechst and in the indicated experiments with a modified lineage cocktail including CD3, B220, CD11b and Gr1 stained with streptavidin. In the indicated experiments, cells were also stained with PE/Cy7-Kit (Abcam) and Epor primary antibody (R&D) and then stained with secondary, anti-goat Cy3 antibody (Jackson Labs).

### Stem and progenitors cell assays

Primitive myeloid progenitors were enumerated using the day 12 CFU-S (CFU-S_12_) assay. Recipient mice were sub-lethally irradiated (7 Gy) using a ^137^Cs-γ-ray source and injected with 10^5^ BM cells. Spleens were isolated 12 days after transplantation and fixed in Bouin’s solution. The number of macroscopic spleen colonies were counted and expressed as the number of CFU-S colonies per 10^5^ donor cells. Clonogenic myeloid progenitors were assessed by standard methylcellulose CFC assays (MethoCult GF M3434; Stem Cell Technologies) using 1.5 × 10^4^ BM mononuclear cells (BMMCs) per well (6-well plate). Colonies were scored after 7 days of incubation and expressed as the number of CFUs per 1.5 × 10^4^ BMMCs before normalizing to total leg (femur/tibia) cell counts.

### HSC transplantation

Recipient mice were irradiated with 9 Gy using a ^137^Cs-γ-ray source. 3–4 hours after irradiation, whole bone marrow cells (marrow cells) from donor animals were infused via the tail vein of recipient animals. Competitive repopulation was used to assess the relative number of HSCs in WBM from *TGFβ1*^ΔMk/ΔMk^ and *TGFβ1*^FL/FL^ mice. *TGFβ1*^ΔMk/ΔMk^ or *TGFβ1*^FL/FL^ WBMCs were mixed with an equal number of marrow cells from congenic CD45.1^+/+^ control mice. The 1:1 (test/competitor) mixture (2_X_10^6^ cells total) was injected into lethally irradiated mice via tail vein. Peripheral blood was analyzed for lymphoid (CD3/CD19) and myeloid (Gr1/CD11b) engraftment by flow cytometry at various times after transplantation. Animals were sacrificed after 16 weeks and marrow harvested for immunophenotyping of HSPC populations and for serial transplantation. Serial transplantation of secondary recipients was performed as for primary recipients except the WBM donor cells were not mixed with additional congenic competitor cells.

### Cell count analysis

To assess blood count recovery, we collected peripheral blood (50 µl) into EDTA-coated capillary tubes (Thermo Fisher Scientific). Differential blood counts were measured using an automated ADVIA 120 Multispecies Hematology Analyzer (Bayer HealthCare) calibrated for murine blood.

### Apoptosis analysis by flow cytometry

Apoptosis in mononuclear cell suspensions was assessed using the Annexin V-FITC Apoptosis Kit (BioLegend, 640914) according to manufacturer’s instructions and analyzed via flow cytometry. In some experiments, apoptosis was assessed using CellEvent™ Caspase-3/7 Green Detection Reagent by incubating marrow cells for 10 minutes prior to analysis by flow cytometry. Cleaved Caspase3 was assessed in surface stained mononuclear cells suspensions permeabilized using CytoFix/Perm (BD). After permeabilization, cells were stained with cleaved-Caspase3 primary antibody (Cell Signaling Technology), washed and then stained with secondary, anti-rabbit antibody Cy3 (Jackson Labs). The cells were then analyzed by flow cytometry.

### Immunofluorescence

Femurs were fixed in 4% PFA overnight, and then decalcified using 10% EDTA before freezing in OCT (Sakura Finetek). Immunofluorescence staining was performed on frozen sections. After blocking with blocking buffer (3% BSA, 3% Serum and 0.03% tween), sections were incubated in primary antibodies overnight at 4°C using anti-cleaved Caspase3 (Cell Signaling Technology) antibody. After washing, sections were stained with secondary antibody (anti-rabbit) CY3 (Jackson Labs). The specificity of staining was confirmed in sequential sections using the secondary antibody alone. Images were acquired using a Zeiss spinning disk confocal microscope or a Zeiss 710 laser scanning confocal microscope.

### In vivo stimulation of erythropoiesis

Human recombinant EPO (300 U/kg body weight, ∼9U per mouse) was injected subcutaneously for 5 consecutive days. On day 6, blood was sampled and mice were killed for analysis.

To induce hemolysis, mice were injected intraperitoneally with phenylhydrazine hydrochloride (PHZ) at 60 mg/kg (Sigma-Aldrich) on day 1 and 2. Blood was sampled on days 3, 5, 8 and 14 from staggered cohorts of mice.

To assess the ability of exogenous TGFβ1 to rescue megakaryocytic TGFβ1 knockout, mice were treated with TGFβ1 (5 μg/kg/day) subcutaneously for 5 consecutive days as indicated. On day 6, blood was sampled and mice were killed for analysis.

### Immunoblotting

Mononuclear cell suspensions were washed in PBS, pelleted and were lysed in lysis buffer (50 mM Tris-HCl (pH 7.5), 120 mM NaCl, and 0.4% NP-40 supplemented with proteinase inhibitors. Samples were transferred to polyvinylidene difluoride (PVDF) membranes (EMD Millipore) and blocked with 5% nonfat dried milk in PBS with 0.1% Tween-20. Primary and secondary antibodies were diluted in blocking solution. Primary antibodies against cleaved Caspase3 (Cell Signaling Technology) was used. Secondary peroxidase-conjugated anti–rabbit antibody (EMD Millipore) was used before chemiluminescent visualization using the SuperSignal West Femto Substrate (Thermo Fisher Scientific).

### Statistical analysis

All data are expressed as mean ± SEM. Student’s *t* test (two-tailed) was used to analyze the statistical differences between groups, with the p-values indicated in the plots (* P < 0.05, ** P < 0.01, *** P < 0.001, or if not shown, the comparison was not significant).

## References

1. Higgins, J. M. Red blood cell population dynamics. Clin Lab Med 35, 43–57, doi:10.1016/j.cll.2014.10.002 (2015).

2. Kassebaum, N. J. et al. A systematic analysis of global anemia burden from 1990 to 2010. Blood 123, 615–624, doi:10.1182/blood-2013-06-508325 (2014).

3. Le, C. H. The Prevalence of Anemia and Moderate-Severe Anemia in the US Population (NHANES 2003-2012). PLoS One 11, e0166635, doi:10.1371/journal.pone.0166635 (2016).

4. Sankaran, V. G. & Weiss, M. J. Anemia: progress in molecular mechanisms and therapies. Nat Med 21, 221–230, doi:10.1038/nm.3814 (2015).

5. Huang, J. & Tefferi, A. Erythropoiesis stimulating agents have limited therapeutic activity in transfusion-dependent patients with primary myelofibrosis regardless of serum erythropoietin level. Eur J Haematol 83, 154–155, doi:10.1111/j.1600-0609.2009.01266.x (2009).

6. Wu, H., Liu, X., Jaenisch, R. & Lodish, H. F. Generation of committed erythroid BFU-E and CFU-E progenitors does not require erythropoietin or the erythropoietin receptor. Cell 83, 59–67 (1995).

7. Hattangadi, S. M., Wong, P., Zhang, L., Flygare, J. & Lodish, H. F. From stem cell to red cell: regulation of erythropoiesis at multiple levels by multiple proteins, RNAs, and chromatin modifications. Blood 118, 6258–6268, doi:10.1182/blood-2011-07-356006 (2011).

8. Malik, J., Kim, A. R., Tyre, K. A., Cherukuri, A. R. & Palis, J. Erythropoietin critically regulates the terminal maturation of murine and human primitive erythroblasts. Haematologica 98, 1778–1787, doi:10.3324/haematol.2013.087361 (2013).

9. Socolovsky, M. et al. Ineffective erythropoiesis in Stat5a(-/-)5b(-/-) mice due to decreased survival of early erythroblasts. Blood 98, 3261–3273 (2001).

10. Ingley, E., Tilbrook, P. A. & Klinken, S. P. New insights into the regulation of erythroid cells. IUBMB Life 56, 177–184, doi:10.1080/15216540410001703956 (2004).

11. Wojchowski, D. M. et al. Erythropoietin-dependent erythropoiesis: New insights and questions. Blood Cells Mol Dis 36, 232–238, doi:10.1016/j.bcmd.2006.01.007 (2006).

12. Menon, M. P. et al. Signals for stress erythropoiesis are integrated via an erythropoietin receptor-phosphotyrosine-343-Stat5 axis. J Clin Invest 116, 683–694, doi:10.1172/JCI25227 (2006).

13. Menon, M. P., Fang, J. & Wojchowski, D. M. Core erythropoietin receptor signals for late erythroblast development. Blood 107, 2662–2672, doi:10.1182/blood-2005-02-0684 (2006).

14. Levesque, J. P. & Winkler, I. G. Hierarchy of immature hematopoietic cells related to blood flow and niche. Curr Opin Hematol 18, 220–225, doi:10.1097/MOH.0b013e3283475fe7 (2011).

15. Wong, P. et al. Gene induction and repression during terminal erythropoiesis are mediated by distinct epigenetic changes. Blood 118, e128–138, doi:10.1182/blood-2011-03-341404 (2011).

16. Zhao, M. et al. Megakaryocytes maintain homeostatic quiescence and promote post-injury regeneration of hematopoietic stem cells. Nat Med 20, 1321–1326, doi:10.1038/nm.3706 (2014).

17. Bruns, I. et al. Megakaryocytes regulate hematopoietic stem cell quiescence through CXCL4 secretion. Nat Med 20, 1315–1320, doi:10.1038/nm.3707 (2014).

18. Lieschke, G. J. et al. Mice lacking granulocyte colony-stimulating factor have chronic neutropenia, granulocyte and macrophage progenitor cell deficiency, and impaired neutrophil mobilization. Blood 84, 1737–1746 (1994).

19. Mori, Y., Chen, J. Y., Pluvinage, J. V., Seita, J. & Weissman, I. L. Prospective isolation of human erythroid lineage-committed progenitors. Proc Natl Acad Sci U S A 112, 9638–9643, doi:10.1073/pnas.1512076112 (2015).

20. Hu, J. et al. Isolation and functional characterization of human erythroblasts at distinct stages: implications for understanding of normal and disordered erythropoiesis in vivo. Blood 121, 3246–3253, doi:10.1182/blood-2013-01-476390 (2013).

21. Chen, K. et al. Resolving the distinct stages in erythroid differentiation based on dynamic changes in membrane protein expression during erythropoiesis. Proc Natl Acad Sci U S A 106, 17413–17418, doi:10.1073/pnas.0909296106 (2009).

22. Dussiot, M. et al. An activin receptor IIA ligand trap corrects ineffective erythropoiesis in beta-thalassemia. Nat Med 20, 398–407, doi:10.1038/nm.3468 (2014).

23. Fenaux, P., Kiladjian, J. J. & Platzbecker, U. Luspatercept for the treatment of anemia in myelodysplastic syndromes and primary myelofibrosis. Blood 133, 790–794, doi:10.1182/blood-2018-11-876888 (2019).

24. Piga, A. et al. Luspatercept improves hemoglobin levels and blood transfusion requirements in a study of patients with beta-thalassemia. Blood, doi:10.1182/blood-2018-10-879247 (2019).

25. Platzbecker, U. et al. Luspatercept for the treatment of anaemia in patients with lower-risk myelodysplastic syndromes (PACE-MDS): a multicentre, open-label phase 2 dose-finding study with long-term extension study. Lancet Oncol 18, 1338–1347, doi:10.1016/S1470-2045(17)30615-0 (2017).

26. Suragani, R. N. et al. Transforming growth factor-beta superfamily ligand trap ACE-536 corrects anemia by promoting late-stage erythropoiesis. Nat Med 20, 408–414, doi:10.1038/nm.3512 (2014).

27. Sanjuan-Pla, A. et al. Platelet-biased stem cells reside at the apex of the haematopoietic stem-cell hierarchy. Nature 502, 232–236, doi:10.1038/nature12495 (2013).

28. Tiedt, R., Schomber, T., Hao-Shen, H. & Skoda, R. C. Pf4-Cre transgenic mice allow the generation of lineage-restricted gene knockouts for studying megakaryocyte and platelet function in vivo. Blood 109, 1503–1506, doi:10.1182/blood-2006-04-020362 (2007).

29. Azhar, M. et al. Generation of mice with a conditional allele for transforming growth factor beta 1 gene. Genesis 47, 423–431, doi:10.1002/dvg.20516 (2009).

30. Pronk, C. J. H. et al. Elucidation of the phenotypic, functional, and molecular topography of a myeloerythroid progenitor cell hierarchy. Cell Stem Cell 1, 428–442, doi:10.1016/j.stem.2007.07.005 (2007).

