## Supplementary figures and images for "Megakaryocyte TGFβ1 Partitions Hematopoiesis into Immature Progenitor/Stem Cells and Maturing Precursors"

### Supplementary-Figure-1

#### Supplementary Figure 1

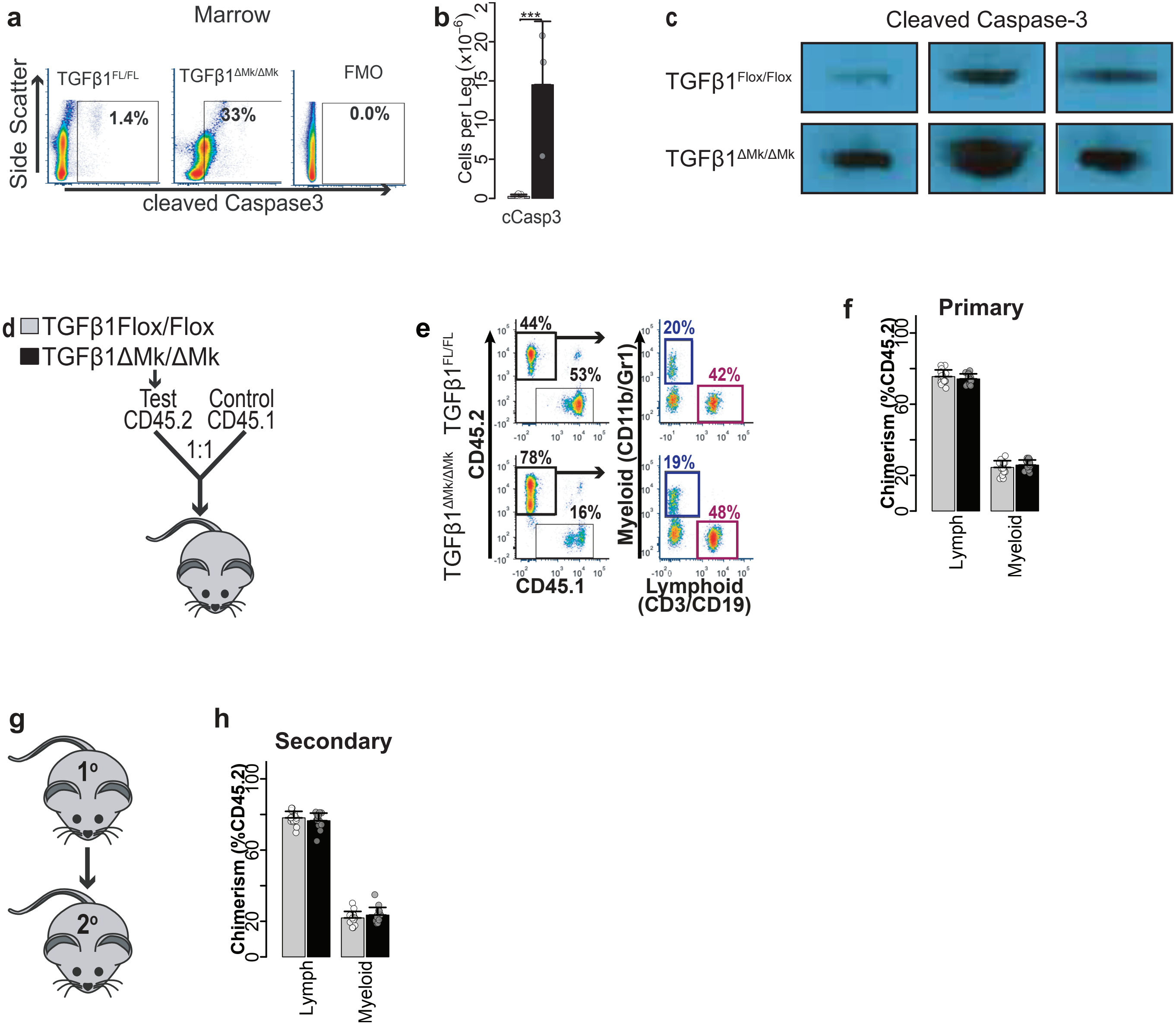

### Supplementary-Figure-2

#### Supplementary Figure 2

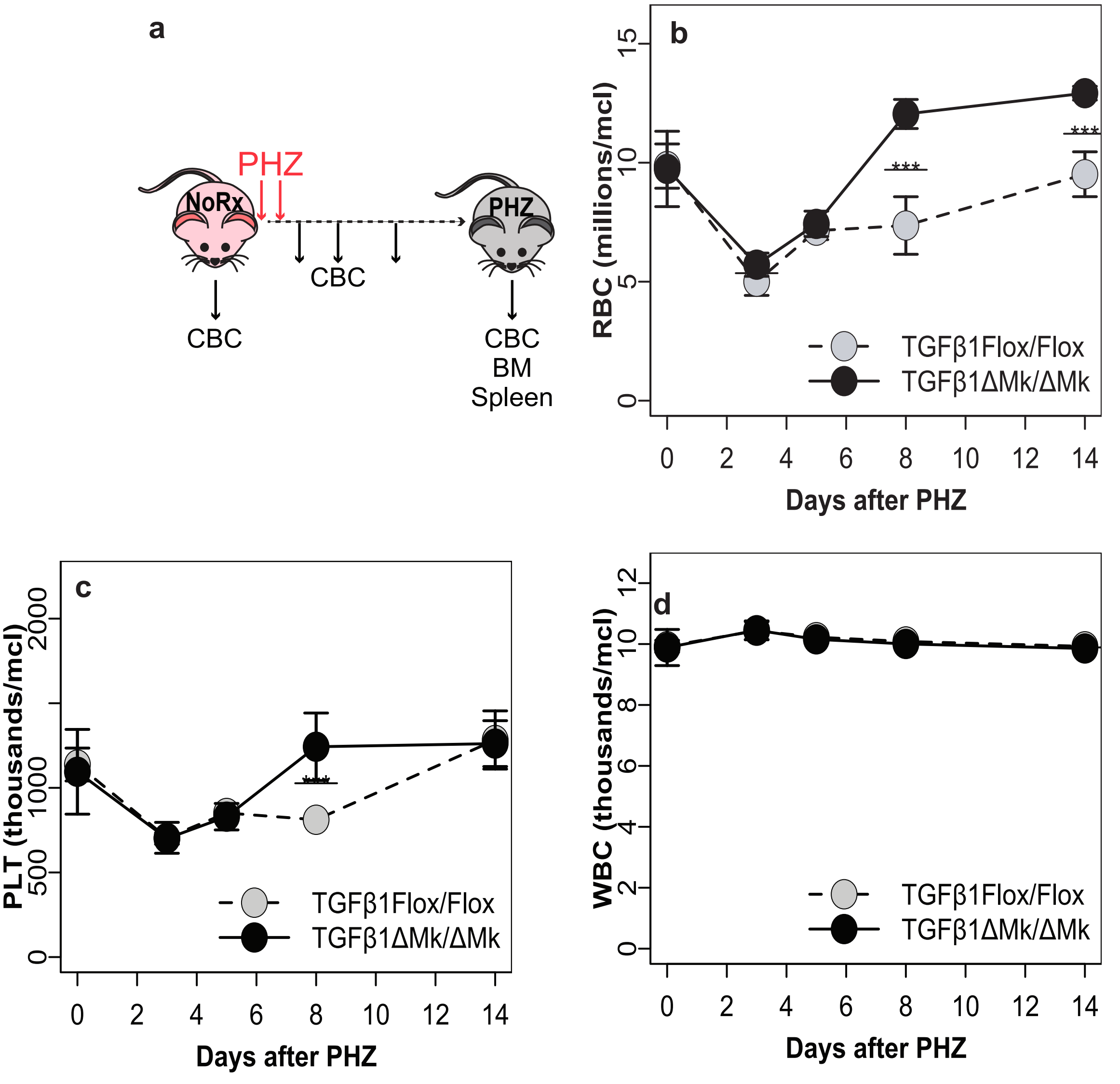

### Supplementary-Figure-3

#### Supplementary Figure 3

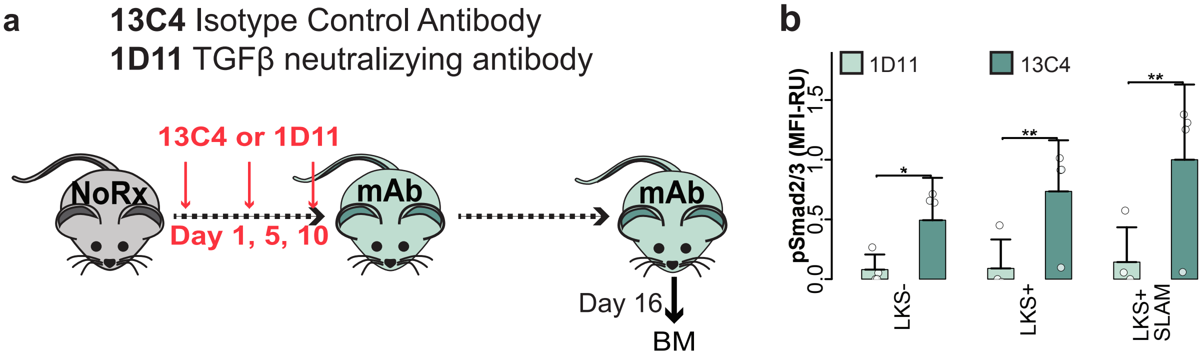
